# Assessing the intrageneric environmental boundaries of the extremophilic cyanobacterium *Chroococcidiopsis* (*Pleurocapsales*) and its implications for space exploration

**DOI:** 10.1101/2022.08.09.503413

**Authors:** John R. Cumbers, Lynn J. Rothschild

## Abstract

The cyanobacterium *Chroococcidiopsis* contains species found in extreme environments, thus providing the opportunity to study adaptation to a range of environments within the confines of a single genus. Due to its extremophilic nature, it has attracted attention for space settlement as well as a model for life elsewhere. In this study, eight unialgal strains from diverse habitats, isolated in unialgal culture and grown in laboratory conditions, were characterized for their ability to survive a range of extreme environments including UVC (254 nm) radiation, oxidative damage, desiccation, and repeated freeze/thawing. The study revealed two previously uncharacterized saltwater isolates of *Chroococcidiopsis* that were more radiation resistant than most of the other isolates. Isolate CCMP 1991 from Hawaii survived up to 1750 J·m^-2^, and isolate CCMP 3184 from Samoa survived up to 1000 J·m^-2^ (254 nm UVR) compared with 250 J·m^-2^ for most other isolates tested. These two UV radiation-resistant isolates are closely related phylogenetically, but inhabit different environments. Each was further characterized for its ability to repair DNA damage as assessed by the repair of UV-induced thymine dimers and for oxidative damage tolerance via resistance to H_2_O_2_-induced (oxidative) damage. Both isolates repaired thymine dimers faster in the light than in the dark with the Hawaiian isolate repairing faster than the Samoan isolate in the light, suggesting repair by photoreactivation. The Hawaiian isolate was more tolerant to H_2_O_2_ exposure than the Samoan isolate, indicating a possible role for antioxidants in the protection of the cell. Both isolates were more tolerant than the other isolates tested to freeze/thawing in liquid nitrogen, which is also known to cause DNA damage. Spectral absorbance scans were performed to detect pigments in each isolate. While all showed peaks likely to be chlorophyll a, carotenoids, phycocyanin, scytonemin and MAAs, the Hawaiian isolate contained a pigment that absorbed at around 325 nm that none of the other isolates contained. Although this pigment is outside the range of UVC absorbance, it is hypothesized that it may play a role in DNA protection as a UV sunscreen or as an antioxidant. The phenotypic similarities in radiation resistance and freeze/thawing resistance among the Hawaiian, Somoan and Negev isolates appear to be the result of environmental adaptation rather than phylogenetic markers as the first of these have been previously shown to be part of a saltwater clade, while the Negev strain falls within a freshwater clade. As pigmentation, and likely resistance to other environmental conditions, can be induced, these data provide a baseline study of strains in an uninduced state. Thus, the true environmental limits to *Chroococcidiopsis* likely go beyond our current knowledge. The implications of this is discussed in relation to space exploration

## Introduction

*Chroococcidiopsis* is a genus of photosynthetic cyanobacteria which has long had a reputation for the ability to survive various extreme environments. As such, the genus is of interest to understand the limits of life on Earth and the evolution of the ability to survive one or more extreme environments (Capece et al., 2013). With these abilities, such organisms make excellent model systems for the search for life elsewhere (e.g., Rothschild & Mancinelli, 2001). Imre Friedmann (1995) suggested that *Chroococcidiopsis* would be suitable for growth on Mars as we move towards human exploration and settlement. They have promise for use in making extraterrestrial bodies habitable (Verseux et al., 2015; Billi et al., 2016), for example, by aiding in the production of desert crusts to mitigate dust (Liu et al., 2008). The genetic capabilities of such organisms could provide these capabilities to other organisms of use that may be more space hardy or genetically tractable (Rothschild, 2010).

Members of the genus have been isolated from diverse environments (Fewer et al., 2002), ranging from the frigid Dry Valleys of Antarctica (Friedmann and Ocampo, 1976) to hot deserts such as the Atacama in Chile (Warren-Rhodes et al., 2006; 2007) and the Negev in Israel (Friedmann et al., 1967). These strains have acquired various traits that allow them to live in their respective extreme environments, such as efficient DNA repair mechanisms to survive in high UV radiation (UVR) environments (Billi et al., 2000a), and desiccation tolerance to survive in low water environments (Caiola et al., 1996a).

Solar radiation is a natural environmental variable for organisms that live on the surface of a planetary body, and exposure to radiation between 400-700 nm is obligatory for photosynthesis. Thus, organisms have evolved in direct response to solar radiation. In contrast, the UV radiation (UVR) portion of the spectrum (10-400 nm) is known to cause a range of different biological stresses (reviewed in Franklin and Forster, 1997). UVR damages biomolecules such as DNA and proteins, which have peak absorption at approximately 260 and 280 nm, respectively. Direct damage from UVR in the UVB (280-315 nm) and UVC (100-280 nm) induces thymine dimer formation, where two adjacent thymine residues are fused together on a strand of DNA (Goodsell, 2001). Several DNA repair mechanisms have evolved to reverse the effects of direct UVR-induced damage, including nucleotide excision repair, base excision repair and photoreactivation (Ehling-Schulz and Scherer, 1999; Friedberg et al., 2005). For example, photolyase enzymes are light-activated DNA repair enzymes, known to work by recognizing cyclobutane-type pyrimidine dimers, and converting them back into monomers in the presence of blue/violet light (Ehling-Schulz and Scherer, 1999; Frohnmeyer, 2003; Faraji and Dreuw, 2014).

UVR also damages other biological molecules in a variety of ways (rev in Rothschild and Cockell, 1999; Sinha and Häder, 2002). UVR can inhibit growth and enzymatic activity (Kumar et al., 1996; Wu et al., 2005), and is known to cause damage to cell membranes, pigments, and signal transduction pathways (Vincent and Roy, 1993; Häder et al., 2003; Cadoret, 2005; Portwich and Garcia-Pichel, 2007).

*Chroococcidiopsis* has been exposed to solar ultraviolet radiation (UVR) in the open environment (Cockell et al., 2008), and simulated Martian UVR environment in the laboratory (Cockell et al., 2005; Billi et al., 2011). Cockell et al. (2005) exposed dried monolayers of *Chroococcidiopsis* sp. 029 to a simulated Martian surface UVR flux as well as visible radiation, resulting in only 1% viability after 5 minutes, and no viability after 30 minutes. However, a 1 mm layer of Martian regolith simulant sieved to a grain size of <45 μM provided protection, similar to that shown for UVR protection by evaporitic salt crusts (Rothschild, 1990). Further studies by Billi’s lab in preparation for ESA’s BioMEX *(BIOlogy and Mars EXperiment)* flight experiment aboard the International Space Station (ISS), confirmed that mineral analogs provide protection to the integrity of the DNA and photosynthetic pigments, even after exposure to 500 kJ·m^-2^ of polychromatic UVR and space vacuum (10^-4^ Pa), exposures that correspond to conditions during a one-year period in Low Earth Orbit (LEO) (Baqué et al., 2014).

Several studies have been conducted through controlled exposure of *Chroococcidiopsis* to UVC radiation at 254 nm. For example, Baqué et al. (2013a) showed that the DNA of dried biofilms of *Chroococcidiopsis* sp. CCMEE 029 maintained DNA integrity virtually identical to unexposed liquid culture controls after exposures of up to 1.000 J·m^-2^ at 254 nm.

Cellular aggregation is correlated with resistance to some environmental factors, including UVR. For example, a dried biofilm of CCMEE 029 exposed to 10 kJ·m^-2^ of UVC radiation protects the lower cell layers by the top cell layers (Baqué et al., 2013a). The cells in the top layer had damaged photosynthetic pigments as suggested by yellowish autofluorescence, while the lower cell layers maintained a red autofluorescence characteristic of chlorophyll. Liquid cultures withstood exposure to 13 kJ·m^-2^ from a germicidal lamp, which Baqué et al. (2013b) ascribe to multiple factors, including cell aggregation which protects some cells from the lethal effects of radiation.

Some strains of *Chroococcidiopsis* are remarkably resistant to x-ray irradiation (0.01-10 nm), even though unlike UVR, naturally occurring exposure is minimal on Earth. Billi and colleagues (2000) exposed ten different isolates from desert and hypersaline environments to x-rays; all showed between 35% and 80% survival at 2.5 kGy, where a Gy is defined as the absorption of one joule of radiation per kg of matter. Survival was reduced 1 to 2 orders of magnitude when the exposure was doubled, and no cyanobacteria survived exposure at 20 kGy. This is nearly as resistant as the most radiation-resistant organisms known can tolerate. For example, Rainey et al. (2005) were able to recover live *Deinococcus*, *Geodermatophilus*, and *Hymenobacter* from the arid soil of the Sonoran Desert after exposure to doses of 17 to 30 kGy. A subsequent study in Billi’s lab was performed on two strains isolated by Roseli Ocampo-Friedmann: *Chroococcidiopsis* spp. CCMEE 029, a cryptoendolith isolated from the sandstone of the Negev Desert (Israel), and CCMEE 057, isolated from chasmoendolithic growth in granite in the Sinai Desert (Egypt). In the desiccated state, both strains survived x-ray exposures up to 5 kGy without detectable damage to either their DNA or membrane integrity (Verseux et al., 2017). To put this in perspective, the average radiation dose from an abdominal X-ray is 0.7 mGy (0.0007 Gy). However, it should also be noted that the effect is correlated with rate of exposure: whereas 5 Gy is lethal to humans in a short period of time, it may be tolerated over a lifetime.

Thus, comparisons between studies with different dose rates are not equivalent. With reference to potential exposure on Mars, an absorbed dose rate of 76 mGy year^-1^ was measured by the Curiosity rover at the surface of Gale Crater (Hassler et al., 2013).

A byproduct of UVR, reactive oxygen species (ROS), can in itself cause oxidative damage to biomolecules (He and Häder, 2002; Pattanaik et al., 2007). The resulting damage often includes single-strand DNA breaks, and the production of 8-hydroxyguanosine residues (Cooke, 2003). Low levels of x-ray radiation also may cause oxidative damage indirectly (Puthran et al., 2009). Repair mechanisms, such as excision repair, can be used for both direct radiation and oxidative damage (rev. in Rastogi et al., 2010).

One mechanism for resistance to radiation and oxidative damage is the production of UVR-absorbing and antioxidant pigments. Mycosporine-like amino acids (MAAs) are some of the strongest natural compounds that absorb in the UVA and are found in a wide phylogenetic variety of organisms including cyanobacteria, red algae, dinoflagellates, corals, and many marine invertebrates. In addition to MAAs, the UVR-absorbing pigments scytonemin is widely distributed among the cyanobacteria, including *Chroococcidiopsis* where scytonemin is found extracellularly in the sheath (Sinha, 2000; Dillon et al., 2002; Řezanka et al., 2004). Some photopigments such as melanin, bacterioruberin and the carotenoids are also known to act as antioxidants (Geng et al., 2008; Karsten and Garcia-Pichel, 1996; Shahmohammadi et al., 1998; Stahl and Sies, 2003; He and Hader, 2002a; b).

A link between desiccation and radiation resistance was established in the exceptionally radiation-resistant bacterium *Deinococcus radiodurans* (Mattimore and Battista, 1996; Battista, 1997). *Chroococcidiopsis* is known to withstand desiccation, although there is strain-specific variation. Fagliarone et al. (2017) revealed two orders of magnitude difference in the ability of 10 strains of *Chroococcidiopsis* to survive four years of air-dried storage. Differences even occurred among isolates collected from the same area.

The ability of *Chroococcidiopsis* to withstand oxidative damage as assessed by protein oxidation was tested after four years of desiccation (Fagliarone et al., 2017). Strain CCMEE 029 from the Negev desert was resistant to 30 min exposure up to concentrations of 500 mM H_2_O_2_, whereas at the other end of the spectrum, CCMEE 584(M) from the Gobi Desert and *C. thermalis* PCC 7203, which was isolated from a soil sample near Greifswald, East Germany, showed significant protein carbonylation at only 10 mM H_2_O_2_. This tolerance to oxidative damage correlated with resistance to γ -ray exposure (Fagliarone et al., 2017).

Billi (2009b) did not find genomic fragmentation, nor loss of membrane integrity in CCMEE 029 (Negev) Cells that had survived desiccation for four years. Protection could be by the production of antioxidant enzymes such as superoxide dismutase (Caiola et al., 1996b, Billi, 2009b), an enzyme known to reduce oxidative stress, and carotenoids (Baqué et al., 2013b).

Combined with the evidence of DNA repair, it seems that *Chroococcidiopsis* may survive in extreme conditions by both DNA protection by antioxidants and by DNA repair.

*Chroococcidiopsis* is thought to able to withstand high levels of X-ray radiation and desiccation due to an efficient DNA repair mechanism (Billi et al., 2000a), possibly the same mechanism as that which is responsible for UVR resistance (Cockell et al., 2005; Billi, 2009a) as both forms of radiation are known to cause strand breakage in DNA. Double-strand breaks in the genomes of a number of *Chroococcidiopsis* isolates, including 029 (Negev) and 171 (Antarctica), were observed following X-ray irradiation by fragmented DNA on an agarose gel (Billi et al., 2000a). This damage was repaired within 24 hours, demonstrating that an efficient DNA repair system existed. Thus, as in *D. radiodurans*, a link between desiccation resistance and radiation resistance in *Chroococcidiopsis* seems likely, although the mechanism of resistance differs as *D. radiodurans* repairs DNA breaks, while *Chroococcidiopsis* prevents them.

Cyanobacteria were responsible for the rise in atmospheric oxygen levels, allowing for the formation of the stratospheric ozone layer. Because cyanobacteria had to access photosynthetically-active radiation at the time when UVR flux on the surface of the Earth should have been substantially higher, cyanobacteria evolved a variety of UVR-protection mechanisms. These include avoiding damage by physical barriers such as living under minerals (e.g., Baqué et al., 2014), or mitigating (e.g., Bebout and Garcia-Pichel, 1995) or repairing damage through physiological means. For example, planktonic *Chroococcidiopsis* are not as resistant to environmental extremes as those in a dried biofilm, including in low Earth Orbit through the EXPOSE R-2 experiment on ISS (Billi et al., 2019) due to both self-shielding and increased production of extracellular polymers.

With interest in *Chroococcidiopsis* high for its potential in space exploration and as a member of the toolkit for space synthetic biology, this study was undertaken to assess intra-generic phenotypic diversity, and to see if a correlation existed between radiation resistance, photolyase activity and pigmentation. The objective was to characterize previously unstudied isolates and compare them to well-studied isolates (e.g., CCMEE 029 Negev and PCC 7203, *C. thermalis*) in order to find novel mechanisms that allow them to survive in extreme environments on Earth in order to transfer these mechanisms to other organisms for the possible use in extreme environments of space (Rothschild, 2010). Although numerous isolates of *Chroococcidiopsis* have been identified and isolated, only a small number of isolates have been studied simultaneously for their abilities to survive in different extreme conditions, and few comparative studies have been conducted to determine the range of adaptive capabilities and mechanisms that have evolved within the genus.

This study was undertaken with the expectation that it would reveal useful phenotypes that could be exploited for space exploration, either by sending *Chroococcidiopsis* or the genes derived from it into space, or by using it as a model organism in the search for life elsewhere. Thus, the focus was on tolerance to UV radiation, desiccation, freeze/thawing and oxidative damage. The mechanism for resistance was approached by assessing the ability of select strains to repair UV-induced thymine dimers in the light and dark, and by obtaining absorbance spectra for each strain. Only strains that were in unialgal culture were selected as they would be the most likely to be used for future study.

## Results

Eight unialgal cultures of *Chroococcidiopsis*, representing a range of environmental sources (Table 1), were assessed for their environmental tolerances. The data indicate a broad range of environmental preferences that are species or strain specific.

**Table 1:** *Chroococcidiopsis* isolates used in this study

| Species | strain | Growth medium | Location isolated |
| --- | --- | --- | --- |
| *unassigned* | CCMP 1991 | BG11 + 4% sea salt | Hawaii |
| *unassigned* | CCMP 2116 | BG11 + 4% sea salt | Nosy Kumba Island, Madagascar |
| *C. polansiana* | CCMP 1489 | BG11 + 4% sea salt | Fernandina Island, Galapagos station 8 |
| *unassigned* | CCMP 3184 | BG11 + 4% sea salt | Island of Savaii, Western Samoa, Micronesia |
| *unassigned* | PCC 6712 | BG11 | Marin County, California |
| *C. thermalis* | PCC 7203 | BG11 | near Greifswald, East Germany. |
| *unassigned* | CCMEE 029 | BG11 | Makhtesh Ramon, Negev Desert, Israel |
| *unassigned* | CCMEE 171-A789-2 | BG11 | University Valley, McMurdo Dry Valleys,<br>Antarctica |

Abbreviations for culture collections: Provasoli-Guillard National Center for Culture Collection of Marine Phytoplankton (CCMP, Andersen et al., 1997; since 2011 the NCMA, the National Center for Marine Algae and Microbiota), the Pasteur Culture Collection of Cyanobacteria (Rippka et al., 1979), The University of Oregon's Culture Collection of Microorganisms from Extreme Environments (CCMEE) (http://cultures.uoregon.edu) and Sammlung von Algenkulturen (SAG) at the University of Göttingen (Göttingen, Germany).

### UVR tolerance

Radiation tolerance was assessed by irradiation with UVC radiation at 254 nm, a wavelength commonly used as a germicidal source. Dose response curves are plotted in Fig. 1. The cells were spotted on the surface of agar plates prior to irradiation to remove the possibility of attenuation by the culture medium, and to minimize self-shading. The figure shows that a range of UVR tolerances exist among the isolates tested. Two isolates survived similarly large doses of radiation: 1991 (Hawaii) which survived up to 1750 J·m^-2^, and 029 (Negev, Israel) which survived up to 1500 J·m^-2^. Isolate 3184 (Samoa) had moderate radiation resistance with survival up to a dosage of 750 J·m^-2^. Isolate 171 (Antarctica) show some survival at 250 J·m^2^, but no survival by 500 J·m^-2^. The remaining four strains (2116, Madagascar; 1489, Galapagos; 7203, Germany; 6712, California) were killed by the lowest dosage of irradiation used in the experiment, 250 J·m^-2^. *Escherichia coli* (Strain TOP10) was irradiated for comparative purposes, and also did not survive the dosage of 250 J·m^-2^.

**Fig. 1.**
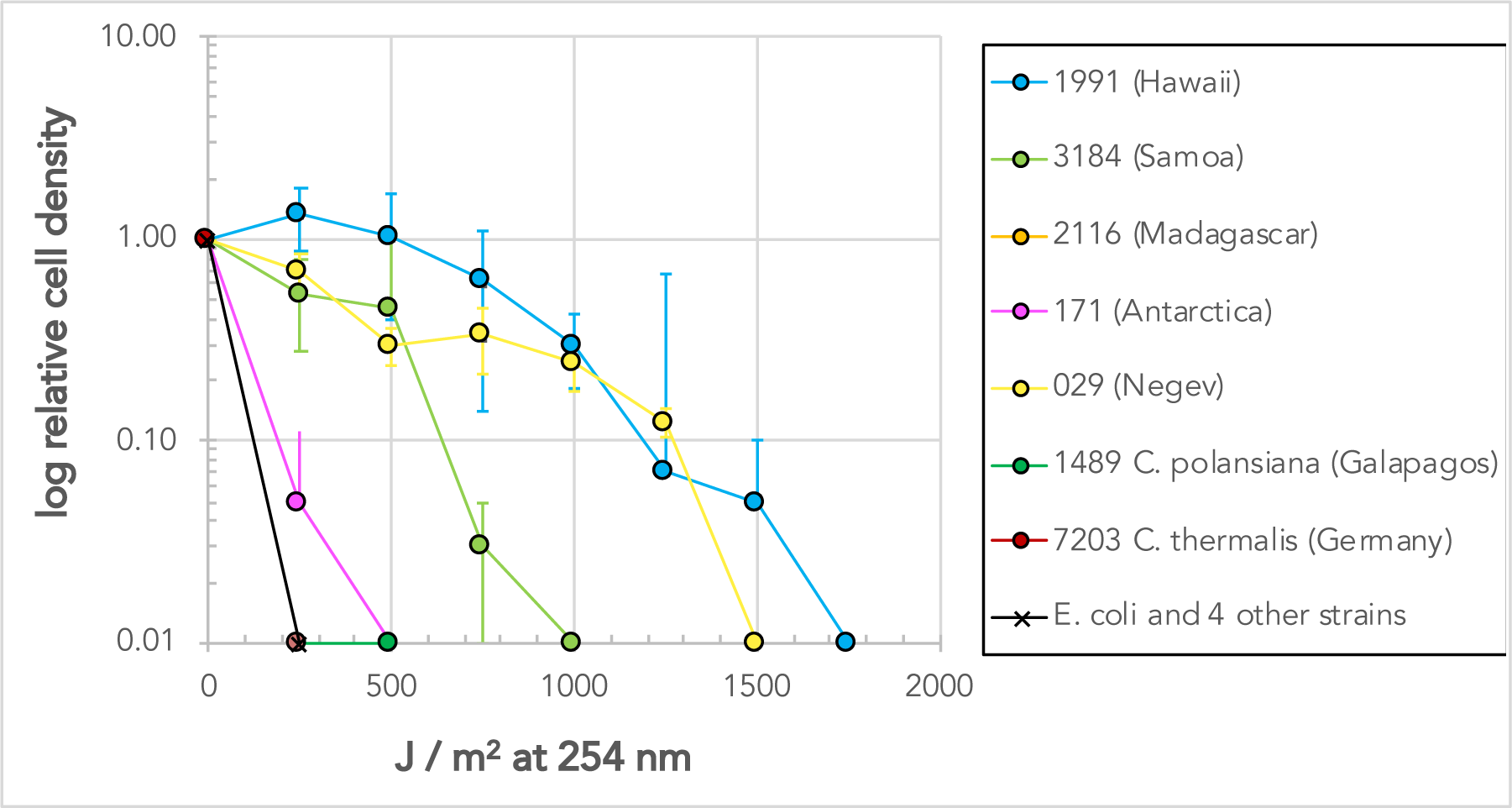
Dose response curves of species exposed to different dosages of UVC (254 nm). Species CCMP 1991 (Hawaii) and CCMEE 029 (Negev) have similar radiation resistance. Species CCMP 3184 (Samoa) has moderate radiation resistance. Cells were irradiated on agar plates and incubated as described above. Cell density was then measured and normalized to the density of un-irradiated cells. Values are means of two independent trials with six replicates per trial.

Previous work has shown that CCMP 1991 (Hawaii) and CCMP 3184 (Samoa) are phylogenetically similar, clustering close to each other in a saltwater clade of *Chroococcidiopsis* (Cumbers and Rothschild, 2014). Because these two isolates are phylogenetically similar but differ in radiation phenotypes, they were investigated in more detail. When triplicate samples of each isolate were exposed to 250 J·m^-2^ of UVC radiation and immediately tested for accumulated damage, CCMP 1991 (Hawaii) contained 73,697 ± 1.7% thymine dimers per Mb of DNA, 8.4% more than CCMP 3184 (Samoa) which contained 67,958 ± 2%. The subsequent repair of the thymine dimers was investigated in both isolates over time as shown in Fig. 2.

**Fig. 2.**
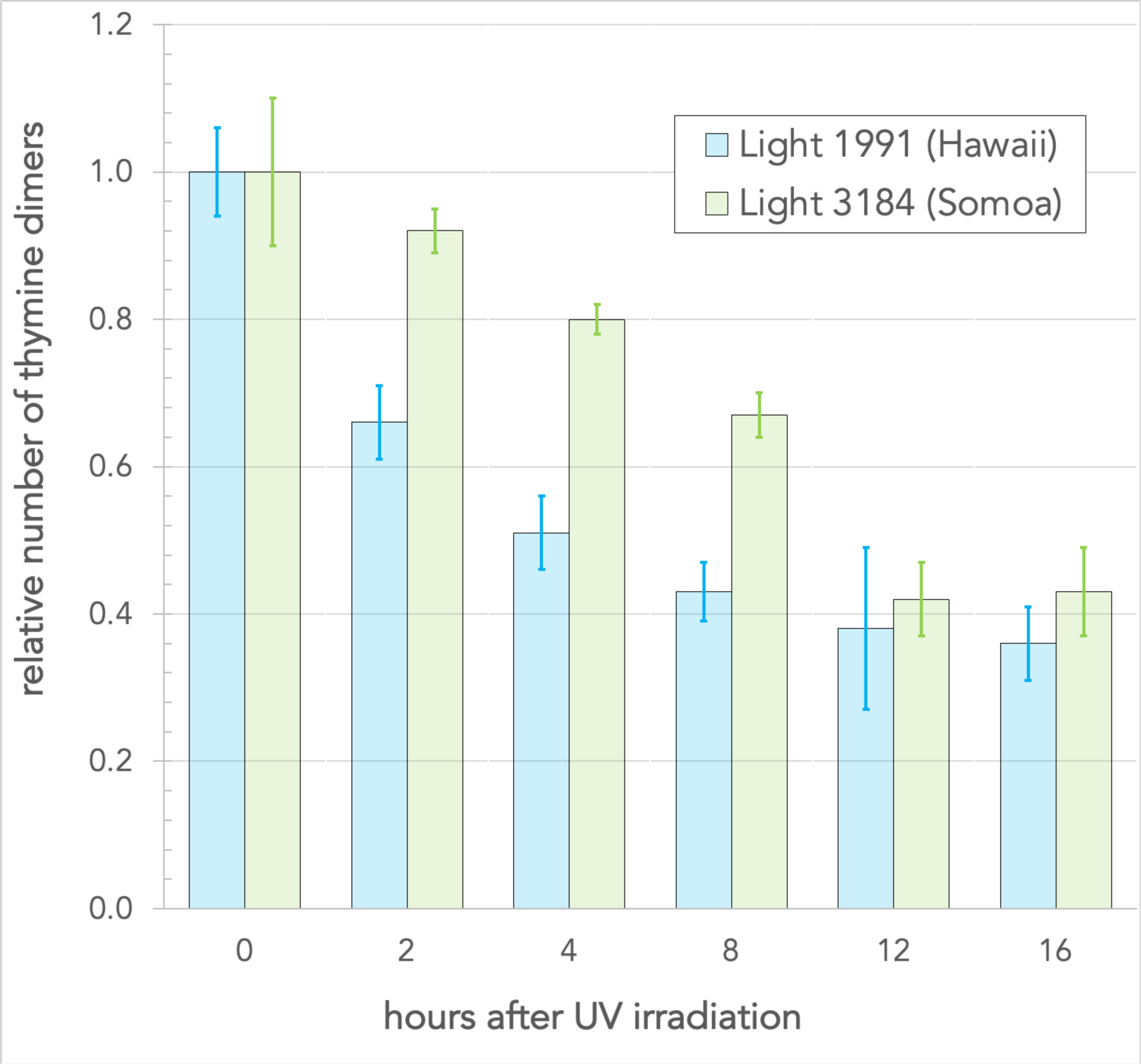
DNA repair of thymine dimers occurs faster in species 1991 (Hawaii) than in species 3184 (Samoa). Cells were irradiated with 250 J·m^2^ of UV radiation and left to recover under laboratory light conditions. Samples were taken periodically during recovery and DNA was extracted. The number of thymine dimers was determined by immunoblotting. Mean values are from three analyses.

Thymine dimer repair occurred faster in in the light in isolate CCMP 1991 (Hawaii) than in isolate CCMP 3184 (Samoa). After four hours, isolate CCMP 1991 (Hawaii) had repaired 51% of the thymine dimers in contrast to only 20% in CCMP 3184 (Samoa). For both isolates, thymine dimer repair takes place faster when cells are left to recover in the light than in the dark (Fig. 3).

**Fig. 3.**
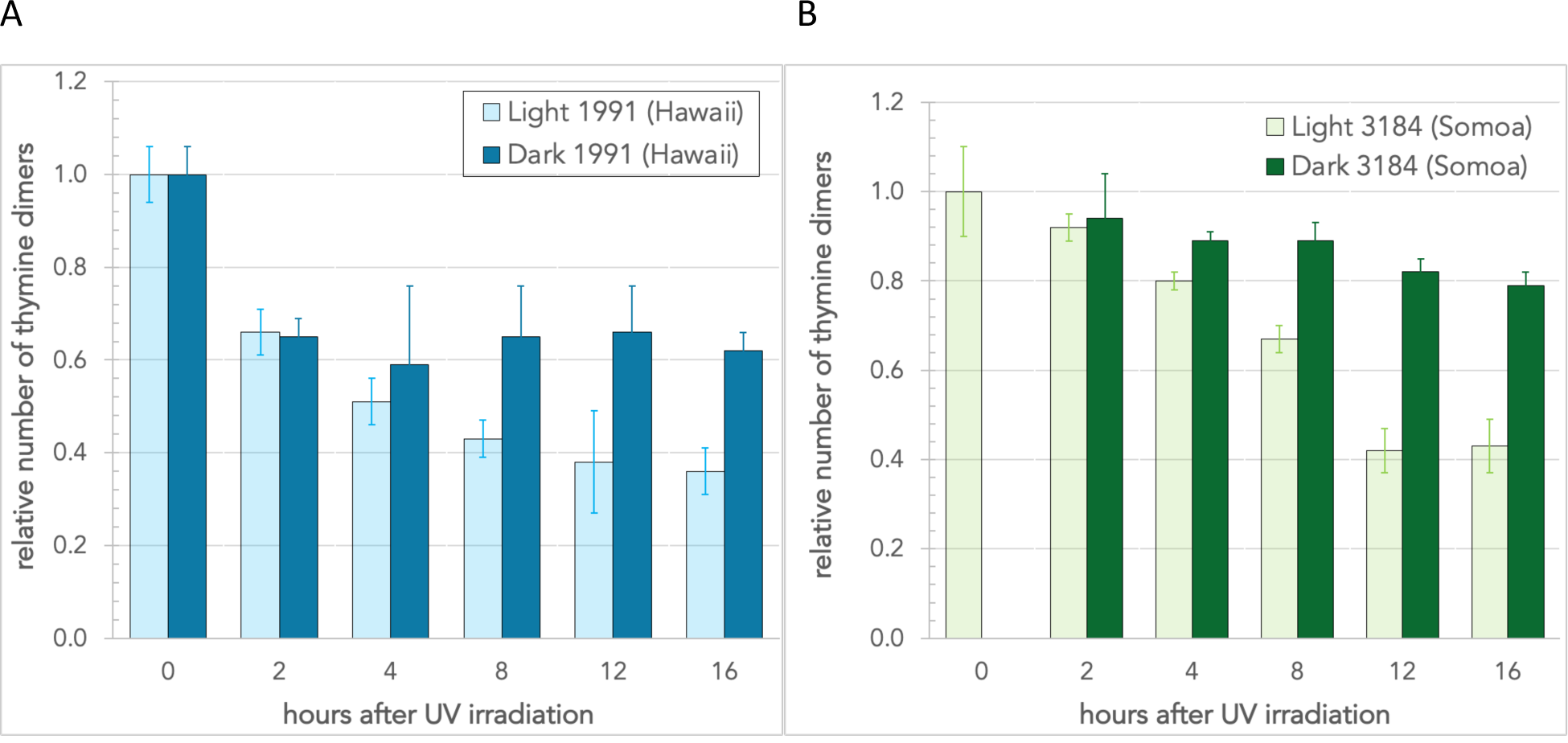
DNA repair takes place faster in the light than in the dark in both 1991 (Hawaii) and 3184 (Samoa). As in Fig. 2, but cells were left to recover in the dark as well as the light. Mean values are from three analyses. A: 1991 (Hawaii); B 3184 (Somoa).

### Oxidative damage tolerance

The ability of the strains to withstand oxidative damage was determined by mixing the cells with varying concentrations of H_2_O_2_, and then plating them on agar plates to determine subsequent colony-forming ability. Isolate 1991 (Hawaii) was more tolerant to H_2_O_2_ than isolate 3184 (Samoa) as shown in Fig. 4. Isolate 1991 (Hawaii) survived up to 100 mM H_2_O_2_, an order of magnitude more than isolate 3184 (Samoa).

**Fig. 4.**
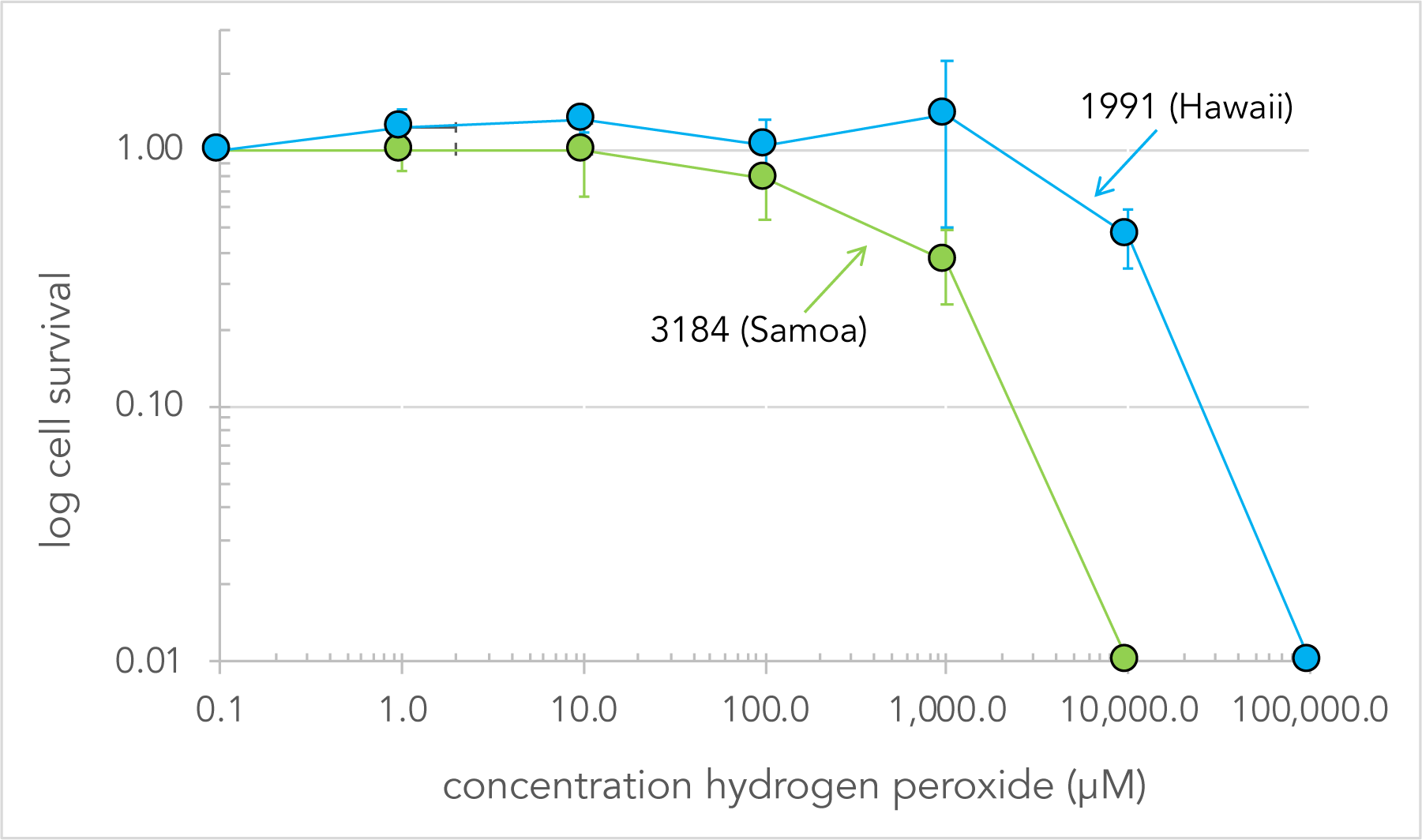
Hydrogen peroxide tolerance in 1991 (Hawaii) and 3184 (Samoa) shows that 1991 (Hawaii) is more tolerant to H_2_O_2_ than 3184 (Samoa). Cells were diluted 1:1 in H_2_O_2_ and plated onto agar plates for growth. Cell density was then measured and normalized relative to the density of un-exposed cells. Mean values are shown from three independent trials with six replicates per trial.

### Desiccation tolerance

Cell survival of air-dried cells after 9 months of storage at room temperature in the dark is shown in Table 2. When the dried cells were re-suspended, only isolate PCC 6712 (California) and CCMP 1489 *C. polansiana* (Galapagos) failed to produce colonies. Thus, under a simple air-drying protocol with storage in the dark, nearly all the strains are desiccation tolerant.

**Table 2:** Colony forming ability after 9 months of desiccation

| Isolate | Desiccation tolerant? |
| --- | --- |
| CCMEE 171 (Antarctica) | + |
| CCMEE 029 (Negev, Israel) | + |
| PCC 7203 (*C. thermalis*) | + |
| PCC 6712 (Marin) | - |
| CCMP 1991 (Hawaii) | + |
| CCMP 1489, *C. polansiana* (Galapagos) | - |
| CCMP 3184 (Samoa) | + |
| CCMP 2116 (Madagascar) | + |

### Freeze/thaw tolerance

Repeated freeze/thawing of cells in liquid nitrogen (Fig. 5a) showed that 1991 (Hawaii) and 3184 (Samoa) both withstood multiple freeze/thaw cycles up to 16 rounds, while the majority of other isolates died after 12 rounds (data not shown.) Isolate 7203 *C. thermalis* died after a single round of freeze/thawing in liquid N_2_. The effect of temperature was assessed by conducting experiments where the cells were frozen in an ethanol/dry ice slurry, which has an approximate temperature of -68 °C, rather than liquid N_2_, which is -196 °C. Isolate 1991 (Hawaii) and 3184 (Samoa) were subjected to freeze/thawing in an ethanol/dry ice slurry (approximately -68 °C). In contrast to the results from freezing in liquid N_2_, neither isolate appeared to be affected by repeated freezing and thawing at the warmer temperate.

**Fig. 5.**
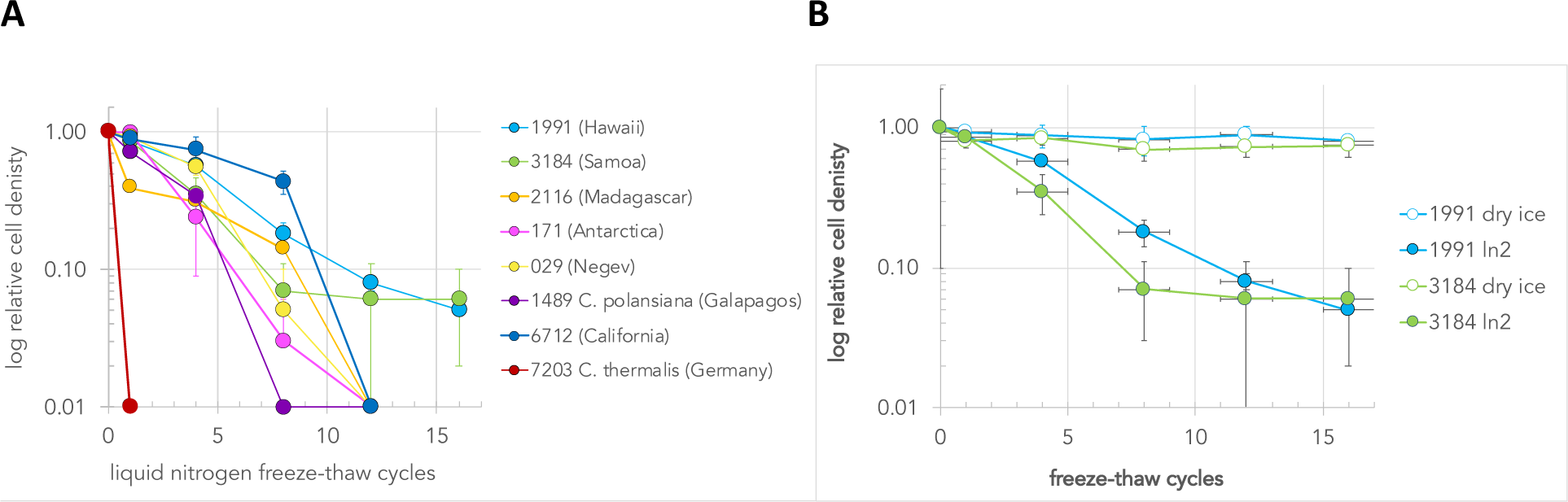
Resistance of strains to repeated cycles of freezing and thawing. Cells were frozen for 30 seconds, and thawed at room temperature. A. Repeated freezing and thawing of cells in liquid nitrogen followed by plating and growth on agar plates. The results show that species 1991 (Hawaii) and 3184 (Samoa) are more resistant to freeze thawing in liquid N_2_ than the other species. Density of cells was assayed and normalized relative to the cell density of control cells. Mean values are shown from two independent trials with six replicates per trial. Species 1489 (Galapagos) and 2116 (Madagascar) are from a single trial with six replicates. B. A comparison of freezing and thawing in liquid N_2_ versus a slurry of dry ice and ethanol. The data are the average of five replicates (1991 dry ice) or six replicates (384 dry ice). Liquid N_2_ data are as in A.

### Pigment absorption

Pigments were extracted in methanol, per standard procedures. The absorbance spectra represent a sampling every 5 nm from 190 to 1000 nm. The spectra show several features expected of cyanobacteria, notably the characteristic peaks of chlorophyll *a* at around 432 and 665 nm (Rowan, 1989, Table 3.2), carotenoids around 490 nm (Erokhina et al., 2002), and phycocyanin at 620 (Sinha et al., 1999; Erokhina et al., 2002). Phycoerythrin (560 nm; Sinha et al., 1999) was not detectable.

Strong absorbance is detected in the UV. The varying strength peaks between 200 and 250 and 325 nm are likely to be the UV-screening pigments scytonemin and mycosporine-like amino acids, based on published studies where the absorption was confirmed by HPLC in the cyanobacterium *Lyngbya* (Sinha et al., 1999, Garcia-Pichel and Castenholz, 1993.) This is especially true of species 1991 (Hawaii) which has a pigment absorbing in the low 300s that is absent in the other species under the growth conditions of this experiment. The identification was not confirmed in this study.

## Discussion

The data presented in this study demonstrate that there is a diversity of phenotypes within the genus *Chroococcidiopsis*. The results show a range of UVR tolerance, indicating that two of the isolates are more tolerant to UVR than the rest. Isolate 1991 (Hawaii) and 029 (Negev) both show similar radiation resistance as seen in Fig. 1. Isolate 029 (Negev) is a well-studied isolate of *Chroococcidiopsis*, and has been shown to be also resistant to X-ray irradiation and desiccation (Billi et al., 2000a) via mechanisms that involve both DNA repair (Billi et al., 2000a) and anti-oxidant production (Caiola et al., 1996b). Isolate 029 (Negev) has also been shown to be UV radiation resistant (Cockell et al., 2005), so it was anticipated that this isolate would shows high radiation resistance in this study too. Here two further isolates *Chroococcidiopsis* that were previously uncharacterized have been identified as UV radiation resistant. Isolate 1991 (Hawaii) is of a similar resistance to 029 (Negev) and isolate 3184 (Samoa) is about half as radiation resistant in comparison.

The results from the radiation exposure, repair and pigmentation experiments presented here, suggest that the two previously uncharacterized isolates survive UV radiation using a combination of DNA protection and repair mechanisms. Isolate 1991 (Hawaii) is more resistant to H_2_O_2_, indicating that it may contain enzymes such as superoxide dismutase (SOD) that play a role in reducing oxidative damage (Caiola et al., 1996b; He and Häder, 2002). Isolate 1991 (Hawaii) is also faster at repairing thymine dimers in its DNA than isolate 3184 (Somoa), indicating that it may have a better DNA repair mechanism.

The large error bars on the dose response curves are due to the clumping of the *Chroococcidiopsis* cells. In general, larger cellular aggregates were observed to contain more cells per aggregate which would likely decrease the chance of genome damage occurring per aggregate, and increase the chance of survivability of at least some members of the colony-forming unit. Isolate 1991 (Hawaii) has tighter packed cells than isolate 3184 (Samoa) which could contribute to the enhanced DNA repair ability observed. *D. radiodurans* is also known to form cellular aggregates which again indicate that this may be part of the DNA protection and/or repair mechanism (Cox and Battista, 2005). So the very characteristic that makes *Chroococcidiopsis* cells difficult to disaggregate may also be the key to its radiation resistance. Billi et al. (2000a) suggested that part of the reason that the genus as a whole may be able to survive such extremes is linked to the morphology of the genus, where mother cells contain daughter cells (baeocytes) at various stages of active cell division. Thus, the fact that the cells were clumping and have baeocytes may contribute to their environmental resistance.

DNA repair was shown to take place faster in the light than in the dark in both 1991 (Hawaii) and 3184 (Somoa), although there was repair in both the light and the dark (Fig. 3). This suggests that a major repair is a light-dependent DNA repair mechanism. Photolyase genes such as the photorepair gene phrA (Hendrischk et al., 2007) are likely responsible for the light repair system, and other mechanisms, such as excision repair, for the dark repair. In particular, isolate CCMP 1991 (Hawaii) appears to contain a faster DNA repair mechanism occurring in the first two hours that is not as apparent in the first two hours of DNA repair in the Samoan (CCMP 3184) isolate. After irradiation, CCMP 1991 (Hawaii) contained more thymine dimers per Mb of DNA than the Samoan isolate CCMP 3184, but it was able to repair these thymine dimers faster, indicating that DNA repair may be more important than DNA protection in this isolate.

The linkage between radiation and desiccation tolerance is known to be through DNA repair (Mattimore and Battista, 1996), as discussed above. However, a link between freeze/thaw tolerance and DNA repair has not been established but could be an important future avenue of research. DNA repair may be a common link between many forms of environmental stresses faced by organisms due to the central role that the molecule plays as an information carrier in the cell. This study focused only on cell survival and DNA damage/repair, and this was limited to thymine dimer damage. Future studies could also investigate other forms of DNA damage that are known to take place (Friedberg, 2005) as well as protein oxidation, which has also been implicated in bacterial radiation resistance (Daly et al., 2007). Fagliarone et al. (2017) demonstrated a clear correlation between desiccation and radiation resistance, and avoidance of protein oxidation. As they point out, mechanistic studies should be conducted examining the possibility of an ROS-scavenging strategy based on a Mn-dependent system such as in *D. radiodurans* (Daly et al., 2004) and for other desiccation- and radiation-tolerant bacteria (Paulino-Lima et al., 2017). Desiccation-resistant organisms usually accumulate protective compounds such trehalose and/or saccharose. Trehalose forms hydrogen bonds with polar residues in proteins and phospholipids in cell membranes and may therefore substitute for water molecules that normally comprises the ‘hydration shell’ surrounding hydrophilic biomolecules, as well as forming a molecular ‘glass’ (vitrification) at biological temperatures (rev. in Morano, 2014). Future mechanistic studies will help explain the higher resistance of desiccated vs hydrated cells to ionizing radiation (Verseux et al. 2017), as well as their endurance when exposed in the dried state to ultraviolet doses simulating 1-year duration in low earth orbit (Billi et al. 2011; Baqué et al. 2013b).

A number of unexpected results exist in the data. Isolate 171 from Antarctica has been previously shown to be resistant to X-ray radiation (Billi et al., 2000a), but in this study, it died after a relatively low UVR dosage of 250 J.m^2^ (Fig. 1). The isolate was also not particularly resistant to freeze/thawing in liquid nitrogen (Fig. 5) when the opposite might be expected from an isolate living in an environment such as Antarctica that undergoes regular freeze/thawing. However, both were resistant to the freeze/thaw regime in the dry ice/ethanol slurry, which is a far closer temperature to that found in the Antarctic (-68 °C, rather than liquid N_2_, which is -196 °C).

The pigment scans in Fig. 6 suggests a pigment in isolate 1991 (Hawaii) that has an absorption peak around 325 nm. This absorption is just in the range of UVA, but distant from 254 nm (UVC), which was the wavelength of the radiation source used in this study. This suggests that this pigment is unlikely to play a role in the protection of the organism from UVC shown here. However, the pigment may still play a role in protection of the organism from UVB. The pigment shows a similar absorbance profile to that of the carotene lycopene (Sicilia et al., 2005), the carotenoid 4-ketomyxol 2ʹ-fucoside (Takaichi, 2005), or a mycosporine-like amino acid (MAA) or scytonemin, both well known from cyanobacteria.

**Fig. 6.**
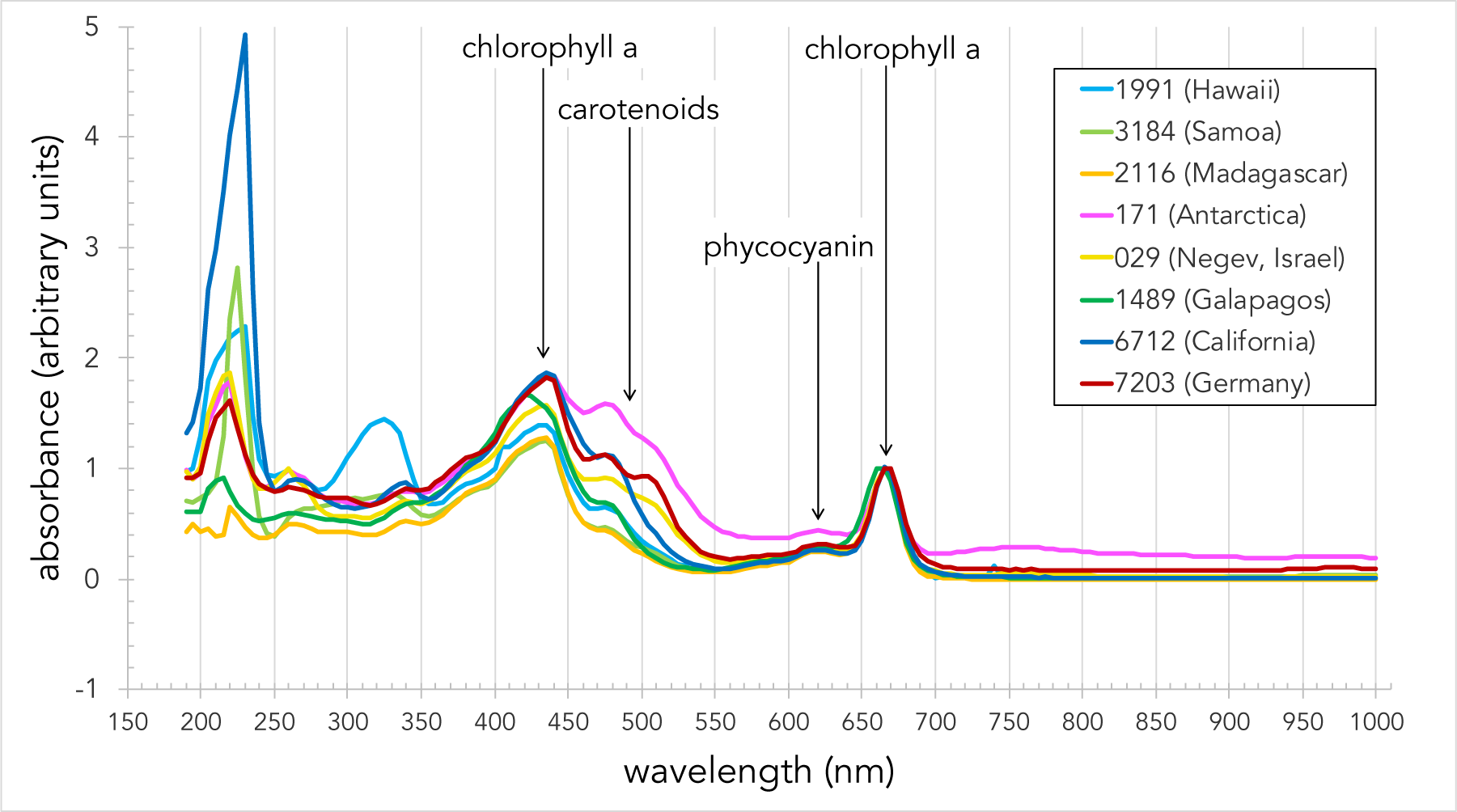
Spectral absorbance scans of pigments extracted in methanol normalized to the red peak of chlorophyll a. The spectra show several features expected of cyanobacteria, notably the characteristic peaks of chlorophyll *a* at around 432 and 665 nm, carotenoids around 490 nm, and phycocyanin at 620. Absorbance in the UV range is likely to be caused by the presence of scytonemin and mycosporine-like amino acids (MAAs). Species 1991 (Hawaii) has a pigment absorbing around 325 nm that is absent in the other species under the growth conditions of this experiment. Identification of the peaks is discussed in the text.

A word of caution should be noted on the interpretation of these data, as well as prior studies with respect to the ability of isolates to withstand environmental extremes. Pigment production responds to the radiation environment, with humans producing more melanin in high UVR environments, and plants changing their pigmentation in response to light levels.

Similarly, the production of MAAs and scytonemin are known to be inducible in cyanobacteria (e.g., Garcia-Pichel and Castenholz, 1993; Ehling-Schulz et al., 1997; Sinha et al., 2003), including in *Chroococcidiopsis* (e.g., Dillon et al., 2002). Thus, the pigmentation profiles in Fig. 6, and many prior studies, may reflect intrinsic pigment production after a number of years in culture in the absence of UVR, rather than the capabilities that these isolates would have shown when collected from their natural environments. The culturing of *Chroococcidiopsis* in the laboratory can, over time, make it less resistant to UV radiation and desiccation (Olsson-Francis and Cockell, 2010a). As the pigmentation is likely affecting the tolerance to UVB irradiation and other stresses, the data presented here should be taken as baseline data from organisms not recently exposed to UVR, and thus provide a pathway to further studies testing environmental resistance and pigmentation after induction.

Based on 16s rRNA and gltX sequence analysis, we suggested previously that the genus *Chroococcidiopsis* contained a saltwater clade within an ancestral freshwater taxon (Cumbers and Rothschild, 2014), and may be polyphyletic. Komárek et al. (2014) have suggested splitting the genus, with the type species, *C. thermalis*, being removed, but the other more extremophilic strains remaining in *Chroococcidiopsis sensu stricto*. The data presented in Fig. 1 might be interpreted as a phylogenetic correlation between UV resistance and the saltwater clade, but the high resistance of the strain from Negev desert (CCMEE 029), which is within the freshwater clade, suggests that this characteristic is an environmental adaptation, and thus polyphyletic. A similar situation exists for the freeze/thaw data presented in Fig. 5, which also shows the Negev phenotypically grouped with members of the saltwater clade, suggesting phenotypic plasticity. Fagliarone et al. (2017) also assessed the relationship between phylogeny and phenotypic characteristics in *Chroococcidiopsis.* These included representatives from deserts worldwide, including the Middle East (Israel, Egypt), Central Asia (Mongolia), South America (Chile) and Antarctica, and the same strain used here to represent the Negev, CCMEE 029, and showed multiple lineages but widespread resistance to desiccation. Even isolates from the same area showed differences in environmental resistance. As Fagliarone et al. (2017) focused on strains from deserts, none of the strains were examined that Cumbers and Rothschild (2014) that they had previously identified as the “saltwater clade” were included.

With baseline data suggesting that 1991 (Hawaii) could be as environmentally robust as 029 (Negev), this strain should be included in future comparisons.

### Implications for space exploration

Cyanobacteria could play an important role in space exploration, from ecopoeisis (making Mars habitable to earthlings) to biological *in situ* resource utilization (ISRU) and human life support (Graham, 2004; Olsson-Francis and Cockell, 2010b; Verseux et al., 2015). Friedmann (1995) first proposed *Chroococcidiopsis* in particular as a suitable organism for terraforming Mars because of its resistance to harsh environmental conditions. While a few terrestrial organisms could probably survive the temperatures and pressures on the surface of Mars (Rothschild, 1990), it unlikely that any terrestrial organism would thrive under those conditions. Even *Chroococcidiopsis* in desert rocks of the Negev need a minimum of −6.9 MPa and 90% RH at 20°C to photosynthesize (Palmer and Friedmann, 1990). Still, the use of polyextremophiles to enable human colonization should result in lower energy expenditures, and thus enhanced mission capabilities as the power is used for other activities.

To test the capabilities of *Chroococcidiopsis* to survive the space environment, the Biofilm Organisms Surfing Space (BOSS) experiment, part of the EXPOSE-R2 space mission exposed cells that were dried as biofilms and compared them to ones dried in the planktonic form (Billi et al., 2019.) Radiation fluences ranged from solar radiation and space vacuum to a Mars-like UVR flux and atmosphere. Upon return to Earth, cells were rehydrated and tested for damage to cell division, the genome and photosynthetic apparatus. Under all conditions, including dark, laboratory-based ground controls, the biofilms recovered better than the planktonic forms, possibly due to the abundance of extracellular polymeric substances, and physiological changes induced by drying. This result confirmed the BOSS team’s prior ground-based studies (Baqué et al., 2013a). While normally biotechnology uses bioreactors as production platforms for microbes, the BOSS results suggest that a biofilm configuration would be a better option for Mars as it would allow the system to better withstand more environmental exposure than planktonic cells, thus potentially lowering infrastructure costs. However, production may be lower in the biofilm than in the planktonic mode, so this would have to be taken into account.

*Chroococcidiopsis* was exposed to space on a second mission on EXPOSE-R2 facility on ISS, BIOMEX (BIOlogy and Mars EXperiment). For this mission, lichens, archaea, bacteria, cyanobacteria, snow/permafrost algae, meristematic black fungi, and bryophytes from alpine and polar habitats were embedded, grown, and cultured on a mixture of martian and lunar regolith analogs or other terrestrial minerals. The organisms and regolith analogs and terrestrial mineral mixtures were then exposed to space and to simulated Mars-like on the EXPOSE-R2 facility. *Chroococcidiopsis* showed survival and recovery on Mars analog minerals, and low DNA-damage particularly in the more protected endolithic dark control areas (de Vera et al., 2019). This result is not dissimilar to prior two-week exposure studies on the free-flying BioPan platform with *Bacillus subtilis* spores where the survival rate was increased by 5 orders of magnitude and more if the spores in the dry layer were directly mixed with powder of clay, rock or meteorites (Horneck et al., 2001). Up to 100% survival was reached in soil mixtures with spores comparable to the natural soil to spore ratio.

While extremophiles for the production of resources in space is of interest to the space community, the addition of synthetic biology greatly expands the potential for biology-based production (Rothschild, 2016). Towards that goal of making *Chroococcidiopsis* a tractable organism for genetic manipulation, Billi et al. (2001) demonstrated gene transformation in several strains of *Chroococcidiopsis* using electroporation and conjugative transfer and showed that CCME 029 was particularly efficient in conjugative transfer. Further, she created two GFP-based plasmids for *Chroococcidiopsis* CCMEE 029 and CCMEE 123. Both were maintained in the host cells after 18 months of dry storage and rehydration. Equally exciting, Billi and colleagues (2000b) transformed *Escherichia coli* BL21DE3 with sucrose-6-phosphate synthase (SpsA) from the cyanbacterium *Synechocystis* sp. strain PCC 6803, conferring a 10,000-fold increase in survival following freeze-drying or air drying. S*ynechocystis* sp. strain PCC 6803 is commonly used in industry and has some tolerance to short-term heat stress, but was isolated from a freshwater lake and has an optimal temperature for growth of 32-38°C (Červený et al., 2015). We propose the exciting possibility of mining the genome of the more extremophilic *Chroococcidiopsis* and using these genes to transform more commonly used synthetic biology chassis organisms such as *Bacillus subtilis* for space use.

This study provides a foundation for performing comparative genomics on isolates of *Chroococcidiopsis* that are phylogenetically similar, but phenotypically diverse, such as CCMP 1991 (Hawaii) and CCMP 3184 (Samoa). These two isolates showed high baseline levels of UV radiation resistance without induction and may prove interesting candidates for future studies in addition to the better studied 029 (Negev). Full genome sequencing or transcriptomics under environmental induction, including UVR, desiccation etc. should illuminate which mechanisms are at play and the relative contributions of DNA protection and repair. Both could lead to a fuller understanding of how *Chroococcidiopsis* survives in extreme environments and how other organisms might one day be engineered to do the same. A confirmation of the identification of UVR-absorbing peaks and their correlation to environmental resistance will allow a fuller understanding of the environmental resistance capabilities of this extremophile genus.

The goal of the study was to begin to understand the mechanisms responsible for survival in extreme environments and to use those as a toolkit to engineering other organisms for survival under such conditions. This was approached by characterizing the response of unialgal isolates. from culture collections to the environmental stresses of UVR, oxidative damage desiccation and freeze/thawing. The results indicated that resistance to UV radiation in uninduced strains is not as widespread in the genus as may have been previously thought. Two isolates were examined in detail for their ability to repair thymine dimers in DNA following radiation and for their ability to survive exposure to H_2_O_2._ Although neither of the mechanisms for these traits has been elucidated, a number of isolates of interest were identified and preliminary work on dissecting the mechanisms of DNA protection and repair were demonstrated. It remains to be seen what the extremophilic capabilities of the genus are after environmental, particularly radiation, induction. The true environmental limits to *Chroococcidiopsis* likely go beyond our current knowledge, making the genus even more interesting for both astrobiology and human space exploration.

## Materials and methods

### Cyanobacterial cultures and culturing

The eight isolates used in this study are listed in Table 1. Each was obtained as a unialgal culture from the Provasoli-Guillard National Center for Culture Collection of Marine Phytoplankton (CCMP; since 2011 the NCMA, the National Center for Marine Algae and Microbiota), the Pasteur Culture Collection of Cyanobacteria, The University of Oregon’s Culture Collection of Microorganisms from Extreme Environments (CCMEE) and Sammlung von Algenkulturen (SAG) at the University of Göttingen (Göttingen, Germany). We are grateful to the late R. Castenholz for the kind gift of isolates from the CCMEE, and to R. Anderson for the kind gift of isolates from the CCMP. The isolates were chosen to represent the geographical, environmental, and phylogenetic diversity of the genus *Chroococcidiopsis* while also being available in unialgal culture. Freshwater isolates were grown in BG11 growth medium (Stanier et al., 1971), made from a 50x concentrate (Sigma-Aldrich Inc., St. Louis, MO, USA, cat. # C3061- 500ML). The pH of the BG11 was adjusted to 7.4 before filter sterilization. Saltwater isolates also were grown in BG11, with the addition of 4% (40 g·L^-1^) sea salt (Sigma-Aldrich Inc., St. Louis, MO, USA cat. # S9883).

Some strains of *Chroococcidiopsis* grow in cellular aggregates making plating of homogeneous monolayers difficult. In order to disaggregate cell aggregates, liquid cultures in mid-exponential phase were first sonicated gently (Ultrasonic processor XL, microtip size 1/16, repeated rounds of three seconds, Misonix Inc, Farmingdale, NY, USA) until the optical density at 730 nm had plateaued, suggesting that large clumps of cells had separated. Cells were then centrifuged at 1000 x g, the supernatant discarded, and the cells resuspended in fresh medium to a concentration of 1·10^6^ colony-forming units mL^-1^ as confirmed by visual inspection with a hemocytometer. Cells were plated on agar plates of the same medium used to grow the isolates, cultured overnight, and inspected the following day to confirm that the sonication had produced a homogeneous monolayer of viable cells. As with previous studies of *Chroococcidiopsis*, a single cell or cell aggregate was counted as a colony forming unit (Billi et al., 2000a). As isolate 1991 (Hawaii) had a morphology showing smaller, denser cell aggregates than 3184 (Samoa), to compare the DNA repair ability of these two isolates, cells were normalized to 6 x 10^7^ cells·mL^-1^ rather than normalizing to colony-forming units.

All growth and survival assays were performed by spotting replicate aliquots of 10 μL of cells onto agar plates, unless otherwise stated in the figure legends. Plates were sealed with parafilm to reduce evaporation before being placed in a growth incubator (Rexmed Inc., Taiwan) at 25°C under a 16h:8h light/dark regime and a mix of Philips TLD36W/840 (cool white) and Philips TLD36w (aqua) fluorescent light at 54±2 μmol photons·m^-2^·s^-1^ as measured with a PAR detector (PMA2100/PMA2132, Solar Light Company, PA, USA). The Model PMA2132 Digital Quantum Light Sensor measures photon flux in the wavelength range 400 to 700 nm.

### Microscopy and image analysis

Agar plates were individually photographed on a light table using a digital camera. Micrographs were taken using a Zeiss Axio Imager Z1 with a DIC filter and a binocular microscope, Zeiss Stemi 2000-c (Carl Zeiss MicroImaging, Thornwood, NY, USA). ImageJ (Abramoff et al., 2004) was used to measure relative cell density. Each agar plate image was cropped to size and converted to a 32 bit grey scale image. The background was subtracted, and light density was measured using the Gel Analyzer tool in ImageJ (Abramoff et al., 2004).

### UVR tolerance

To determine the UVR tolerance of each isolate, cells were first plated onto agar plates in 10 µL spots as described above. The agar plates were then covered with sterile Petri dish lids and left for 24 hours before irradiation. After this time, lids were removed, and cells were placed in a sterile hood 44 cm beneath a UVC light source of 254 nm for different lengths of time to achieve various dosages of radiation. The intensity of the UVC source was 42 µWs·cm^2^ at 254 nm, measured with a high-resolution spectrometer (HR4000, Ocean Optics, Dunedin, FL, USA). Dosages were also measured in real time using a radiometer (PMA2100, Solar Light Company, Glenside, PA, USA). For comparison, *Escherichia coli* (isolate TOP10, Invitrogen, Carlsbad, CA USA) was irradiated after plating cells on LB medium in the same manner as the *Chroococcidiopsis*.

### Thymine dimer detection

Thymine dimers were measured using a dot blot assay according to Sinha et al. (2001) as modified by Leuko et al. (2011). DNA from irradiated cells was extracted using a DNeasy Plant Mini Kit (Qiagen, Gaithersburg, MD, USA) according to the manufacturer’s instructions. The quality (OD 260/280 ratio) and quantity of the extracted DNA was measured on a Nanodrop (ThermoScientific, Wilmington, DE, USA). The DNA was diluted to 2 ng·µL^-1^, NaOH added to 1/10^th^ volume, and incubated at 80° C for 30 min. The denatured DNA was then blotted onto a Hybond LFP membrane using a Minifold I dot blot system (both from GE Healthcare, Piscataway, NJ, USA) followed by a wash with 400 µL TE buffer per well. The membrane was dried for 30 min at 80° C in a drying oven before being blocked with 5% ECL advanced blocking reagent (GE Healthcare, Piscataway, NJ, USA) for one hour. The unwashed membrane was incubated in mouse anti-thymine dimer primary antibody (#MC-062, Kamiya Biomedical, Seattle, WA, USA) at a 1:3000 dilution in PBS-T buffer for two hours at room temperature. The membrane was then washed twice with PBS-T followed by a one hour incubation in a secondary antibody, sheep anti-mouse IgG with Cy3 labeling (Jackson ImmunoResearch Laboratories, Inc., West Grove, PA, cat # 515-165-062) at a 1:1,000 dilution in PBS-T with 5% ECL advanced blocking reagent. The membrane was washed twice in PBS-T before being visualized on a Typhoon Trio Scanner (GE Healthcare, Piscataway, NJ, USA) and quantified with ImageJ (Abramoff et al., 2004).

### Oxidative damage tolerance

To assess the effect of an oxidant on survival, the cells were exposed to hydrogen peroxide. H_2_O_2_ dilutions were freshly prepared from a 9.7 M, 30% v/v stock of H_2_O_2_ (Sigma-Aldrich Inc., St. Louis, MO, USA) in fresh media. Cells were thoroughly mixed 1:1 for each dilution in PCR tubes by pipetting up and down, immediately plated and growth assayed as described above.

### Desiccation tolerance

Desiccation tolerance was assessed in air-dried cultures, kept in the dark at room temperature, according to Billi (2009b) and Caiola et al. (1993). After 9 months, cells were rehydrated by suspension in 50 µL of sterile water rather than fresh medium in order to keep the solute concentration as it was before desiccation. The cells were re-plated, and their growth assayed via microscopy.

### Freeze/thaw tolerance

To measure tolerance to repeated cycles of freezing and thawing, 70 µL of a liquid culture of each isolate was placed into a 0.2 mL PCR tube. The tube was then flash frozen by placing it for 30 sec into an ice bucket containing liquid nitrogen. The tube was then removed and allowed to thaw at room temperature. Cells were then repeatedly re-frozen or plated in 10 µL replicates and growth was assayed as described above. The same protocol was followed for freeze/thawing in the dry ice/ethanol slurry.

### Pigment extraction and absorbance measurements

Pigments were extracted in the dark by first centrifuging 1.5 mL of 1·10^6^ cells·mL^-1^ liquid cultures at 4000 x g for 3 min. The supernatant was removed, leaving behind ∼50 µL of liquid and ∼25 mg of wet weight. Tubes were then placed in ice, and cells were ground with an electric pellet pestle motor (Kontes/Sigma-Aldrich Inc., St. Louis, MO, USA) in repetitions of 30 sec until cells were broken, as confirmed by light microscopy. An aliquot of 250 µL of methanol was added, and the suspension mixed by hand. Cells were centrifuged at 10,000 x g for 10 min. The supernatant containing the extracted pigment was removed, and the absorbance measured from 190 to 1000 nm in 10 nm intervals on a Spectramax 384 Spectrophotometer (Molecular Devices, Sunnyvale, CA, USA).

## Acknowledgments

The laboratory work was performed in partial fulfillment of the following Ph.D. thesis: Cumbers, J.R. Phylogenetic and Phenotypic Studies of the Extremophile *Chroococcidiopsis* (Cyanobacterium). A Foundation for Synthetic Biology in Space. Ph.D. Thesis, Brown University. 2012. https://doi.org/10.7301/Z0VX0DTV. We are enormously grateful to the other members of the thesis committee, Gary Wessel, Mark Tatar, Casey Dunn, and Daniel Weinreich for technical advice and support. We thank S. Pete Worden for funding this study while Center Director of NASA Ames Research Center, and for encouraging LJR to explore the potential for synthetic biology as an enabling technology for NASA’s missions. This work was in part funded by NASA Cooperative Agreement Award Number, NNX08AZ52A.

## Supplementary information

None

## Related Manuscripts

Cumbers, J. & **Rothschild**, L.J., 2014. Salt Tolerance and Polyphyly in the Cyanobacterium *Chroococcidiopsis* (Pleurocapsales). *J. Phycol.* **50**: 472–482.

Ph.D. thesis: Cumbers, J.R. Phylogenetic and Phenotypic Studies of the Extremophile *Chroococcidiopsis* (Cyanobacterium). A Foundation for Synthetic Biology in Space. Ph.D. Thesis, Brown University. 2012. *Molecular Biology, Cell Biology, and Biochemistry Theses and Dissertations.* Brown Digital Repository. Brown University Library. https://doi.org/10.7301/Z0VX0DTV

